# Molecular and Structural Characterisation of the MERS-Coronavirus Polymerase

**DOI:** 10.64898/2026.11.01.763422

**Authors:** Rory F Cunnison, Sonu Subin Philip, Adrian Deng, Danielle Groves, Angus M Spence, Christopher M Coleman, Jeremy Keown, Nicole C Robb

## Abstract

MERS-CoV (Middle East respiratory syndrome coronavirus) belongs to the *Betacoronavirus* genus, which also includes SARS-CoV-2, the causative agent of the COVID-19 pandemic. MERS-CoV is the pathogen responsible for Middle East Respiratory Syndrome (MERS), a respiratory disease with a 36.5% case fatality rate. Given the capacity of coronaviruses for genetic recombination and mutation, MERS-CoV remains a pathogen of significant pandemic potential, with viral evolution potentially facilitating enhanced adaptation and transmission in humans. There are no approved treatments for MERS-CoV infection. The MERS-CoV replication-transcription complex (RTC), a multimeric protein complex key to genome amplification, has no cellular analogues and a highly conserved sequence, making it a useful target for antiviral compounds; however, structural and biochemical characterisation of the MERS-CoV RTC is limited. Here, we characterise the minimal MERS-CoV RTC using in vitro and single-molecule assays and show that it is susceptible to the RNA-dependent-RNA-polymerase inhibitor suramin. We present a structure of the minimal MERS-CoV RTC in complex with an RNA duplex, which shows strong structural similarity to the SARS-CoV-2 RTC. We use cross-subunit swapping experiments to show that the MERS-CoV NSP12 core can efficiently utilise SARS-CoV-2 cofactors NSP7 and NSP8, whereas SARS-CoV-2 NSP12 exhibits reduced in vitro activity with the MERS-CoV NSP8. This asymmetric cross-species compatibility highlights both the evolutionary conservation and virus-specific adaptation of coronavirus polymerase-cofactor interactions. Together, these findings provide a structural and mechanistic framework for understanding MERS-CoV RNA synthesis and establish its conserved RTC as a potential target for the development of broadly acting coronavirus antivirals.

**Significance Statement:** Middle East respiratory syndrome coronavirus (MERS-CoV) causes severe respiratory disease and represents a significant pandemic threat because of the capacity of coronaviruses to mutate and recombine. Despite this risk, there are no approved vaccines or specific treatments for MERS, and no structure of the essential replication-transcription complex (RTC) has been available. Here, we define the biochemical properties of the MERS-CoV RTC and determine its structure bound to RNA, revealing conserved and virus-specific features of coronavirus replication. We further demonstrate inhibition of the polymerase by suramin. By establishing a structural and functional framework for MERS-CoV genome replication, this work addresses a key gap in our preparedness for MERS-CoV and supports the development of antivirals that could be effective against multiple coronaviruses.

## Introduction

Since the turn of the millennium, *betacoronaviruses* have been responsible for a series of severe epidemics and pandemics. The severe acute respiratory syndrome (SARS) outbreak of 2002-2004 resulted in over 8,000 human cases, with a mortality rate of approximately 10% (1). Middle East respiratory syndrome (MERS) emerged in 2012 and remains an ongoing public health threat, with sporadic cases and outbreaks continuing to occur; as of May 2026, 2,637 confirmed cases and 966 deaths have been reported, corresponding to a case fatality rate of 36.5%. (2). Finally, the COVID-19 pandemic, caused by Severe Acute Respiratory Syndrome coronavirus 2 (SARS-CoV-2) and beginning in late 2019, has resulted in over 770 million infections and 7 million deaths to date (2). Other zoonotic *betacoronaviruses* also exist, but infections generally result in mild, flu-like symptoms (3).

The recent and potential threat to life from members of the *Betacoronavirus* genus requires the development of antiviral treatments that are robust enough to navigate the high mutation rates (4) and genomic recombination during co-infection (5), (6) inherent to these viruses. At present, FDA approved direct antiviral treatments for infections of SARS-CoV-2 include Remdesivir, Molnupiravir, and Paxlovid (7). MERS-CoV and SARS-CoV infections currently have no approved antiviral treatments, although Ribavirin and Lopinavir/ritonavir treatments were used during the respective outbreaks, with limited effect (8).

The coronavirus family are so-named due to the crown-like spike glycoproteins which are deposited on the surface of the viral envelope, alongside the membrane and envelope proteins. These proteins, along with nucleoprotein, accessory, and non-structural proteins (NSPs), are encoded in a positive-sense, non-segmented, single stranded genome of ∼30 kDa (9). The size of this viral genome is greater than comparable viruses and is believed to be due to the activity of the proofreading enzymes, such as the NSP14 protein, which can correct the high error rates of the low-fidelity NSP12 RNA-dependent-RNA-polymerase (RdRp) (10). The genome is replicated and transcribed by the dedicated viral RdRp, producing negative strand copies of the genome (gRNA) which are transcribed into positive strand genomes, or subgenomic mRNAs (sgRNA) which are translated into viral proteins (11), (12). Whilst initial translation of the genome occurs directly in the cytoplasm, subsequent rounds of genome replication and transcription are believed to occur in double membrane vesicles (DMVs), reshaped by viral NSPs from the endoplasmic reticulum (13). This approach obscures the production of nascent viral RNA from cellular immune responses.

The SARS-CoV and SARS-CoV-2 RdRps form replication-transcription complexes (RTCs), which can contain helicase, 3’-5’ exoribonuclease, and 2’-O-methyltransferase RNA capping subunits to fulfil both replication and transcription requirements (14), (10), (15). However, the minimal RTC – capable of performing template-specific replication and transcription – is composed of just three NSPs: NSP12, NSP7, and two copies of NSP8 (16), (17). Limited biochemical data suggests that the MERS-CoV RTC is also likely composed of NSP 12, 7 and 8 (18).

In SARS-CoV and SARS-CoV-2 NSP12 contains the core catalytic centre of the RdRp, including highly conserved motifs such as fingers, thumb and palm domains typical of single-subunit RNA polymerases, but also a partially disordered N-terminal *Nidovirus* RdRp-associated nucleotidyl transferase (NiRAN) domain, which catalyses the formation of viral 5’ RNA caps, and an interface domain connecting the N- and C-terminal units (17), (19). NSP7 interacts with both a fingers extension near the thumb of NSP12 and one copy of NSP8. It has been shown to be essential for extension activity in both SARS-CoV and SARS-CoV-2 RTCs (20), (17), (19).

The minimal RTC also contains two copies of NSP8. Cryo-electron microscopy (Cryo-EM) structures of the SARS-CoV-2 RTC show that NSP8 behaves as a scaffold, contacting duplexed RNA up to 28 nucleotides from the RTC active site using positively charged N-terminal domains. The N-terminal domains, along with helix-forming linker residues connecting to the C-terminal domain, form so-called “sliding poles”, which prevent early complex dissociation from the RNA and improve RdRp processivity by contacting the RNA on exit from the active site. The C-terminal domains of the two NSP8s complex with NSP7 and NSP12, with one copy binding to NSP7 and partially to NSP12 near the thumb region (NSP8b). The other copy binds to NSP12 near the finger subdomain, also interacting with the NSP12 interface region (NSP8a) (17), (21), (16), (22). There are multiple structures showing multimeric complexes of NSP8 and NSP7 in the absence of NSP12; the C-terminal domain of NSP8 appears to have a strong affinity for NSP7, producing structures of a hexadecameric complex (23). Furthermore, this hexadecameric complex was proposed to have both primed and de novo extension activity, and NSP8 alone was proposed to have weak “primase” activity (24), (25). How these alternative structures contribute to RTC formation or to replication and transcription of coronavirus gRNA is unknown at present.

Despite the continuing pandemic potential and sporadic outbreaks caused by MERS-CoV, fundamental questions remain regarding the composition, biochemical properties and molecular architecture of its replication machinery, as well as the extent to which RTC structure and function are conserved across divergent *betacoronaviruses*. Here, we address these questions by biochemically and structurally characterising the minimal MERS-CoV RTC, defining its susceptibility to polymerase inhibition, and testing the functional interchangeability of RTC components between MERS-CoV and SARS-CoV-2. We produced a functional recombinant minimal MERS-CoV RTC which displayed primer extension activity in both in vitro and single-molecule assays and produced a cryo-EM structure of the complex which allowed us to compare MERS-CoV with the more well-defined SARS-CoV-2 RTC. Together, these findings establish a structural and functional framework for MERS-CoV RNA replication that will help to inform the future development of antiviral strategies against these viruses.

## Results

### The minimal MERS-CoV RdRp is a multiprotein complex consisting of NSP12, 7, and 8

We initially established a minimal MERS-CoV RTC by expressing and purifying three NSPs believed to form the complex and produce extension activity: NSPs 12, 7, and 8. NSP12-His10-TEV-protein A plasmid constructs were expressed in sf9 insect cells, whilst GST-3C-NSP8 and GST-3C-NSP7 constructs were expressed in *E*.*coli* cells. Both NSP12 and NSP7/8 were purified first through an affinity capture step before additional purificaiotn using size-exclusion chromatography. Denaturing SDS-PAGE analysis of the proteins showed proteins of ∼110 kDa, ∼ 23 kDa, and ∼ 9 kDa for NSP 12, 8, and 7 respectively, equal to the predicted molecular weights of the proteins (Figure 1A). Additional bands in the electrophoresis gels are by-products of the purification process and did not affect characterisation of the proteins.

**Figure 1:**
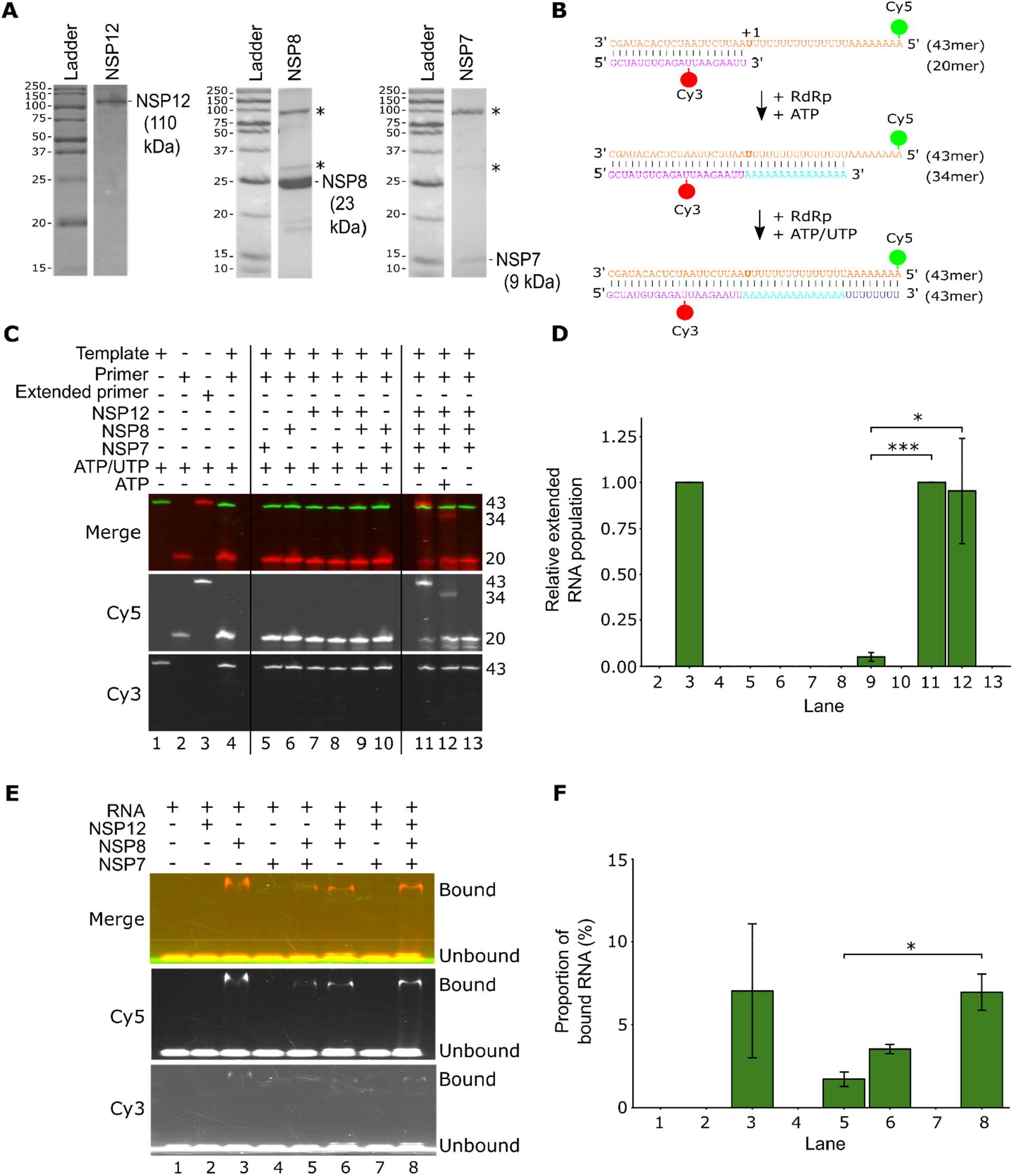
The expressed and purified minimal MERS-CoV RTC is transcriptionally active and binds duplex RNA via NSP8. **A)** SDS-PAGE lanes validating the size of purified MERS NSP12, 8 and 7 proteins. NSP8 and NSP7 preparations had trace contaminants 3C protease (64kDa) and GST-tag (26kDa) (marked by ^*^) but had no effect on proceeding assays. **B)** Sequences of the annealed template (orange) and primer RNA (purple), before and after RdRp-mediated extension. **C)** A representative *in-vitro* extension assay in which individual MERS NSP12, 8 and 7 proteins were assayed in various combinations and the extension products visualised via a urea denaturing gel. Lanes 9, 11 and 12 show extension based on the minimal presence of NTPs, NSP8 and 12, with NSP7 considerably improving extension activity. **D)** Quantification of the *in-vitro* extension assay of MERS RTC. Extension activity is normalised to lane 11 (RNA with NSP12, 8 and 7 in ATP and UTP) with SEM error bars. All conditions have a minimal n=3. NSP12 with NSP8 activity (lane 9) was significantly reduced (P=0.0008 and P=0.044, respectively) compared to conditions with NSP12, 8 and 7 in ATP and UTP (lane 11), and with ATP only (lane 12). Statistical comparisons were conducted using student ‘s unpaired t-test. **E)** A representative EMSA of NSP combinations from 1B. Individual NSP proteins (Lanes 2-4) and protein combinations reveal NSP8 drives RNA binding by the complex regardless of protein partners. Gels were visualised in Cy5 and Cy3 channels and combined to produce a composite. **F)** Quantification plot of EMSA binding assay of MERS RTC. RNA binding was normalised relative to lane 8 (RNA with NSP12, 8 and 7), and plots show the mean proportion of complex bound RNA. Error bars represent the standard error of the mean of a minimum of 2 independent repeats. A student unpaired t-test revealed one significant difference between NSP7+8+12 and NSP7+8 conditions (P= 0.037).

We next used an in vitro fluorescent extension assay to investigate which components of the minimal RTC could extend an RNA primer-template duplex (26). A ratio of 1:5:5 of NSP 12:7:8 was used in our extension assays, consistent with ratios shown to produce high complex activity in similar assays with the minimal SARS-CoV-2 RTC (27). We supplied individual NSPs or preincubated combinations of NSP12, 7 and 8 together before adding annealed fluorescent primer-template duplexes to the reaction mixture and incubating. The template sequence was based upon a 20-nucleotide coronavirus 3’ UTR promoter sequence (28) followed by a tract of 15 Us and 8 As. Upon addition of ATP alongside the RTC, we anticipated extension of the primer by 15nt to produce a 35mer product, whilst addition of ATP and UTP would generate a full-length 43nt product (Figure 1B). The combined individual, pairwise and full RTC NSP reaction mixtures were run on a denaturing electrophoresis gel (Figure 1C&D). As anticipated, addition of individual NSPs did not produce extension products upon addition of ATP and UTP, consistent with assays performed on the SARS-CoV and SARS-CoV-2 RTCs (20), (17), and neither did the addition of NSP12+7 or NSP7+8. However, the addition of NSP12+8 resulted in a small amount of primer extension (∼8%), which increased to 100% when all three NSPs were added together (Figure 1C, lanes 9, 11 and 12), confirming that all three NSPs are required for full transcriptional activity in vitro and highlighting the crucial role of NSP7 in performance of the minimal RTC.

To understand which components of the minimal complex was driving RNA binding, we assayed combinations of NSP7, 8, and 12 with the same fluorescent template-primer duplex under native conditions. Biochemical data from SARS-CoV, SARS-CoV-2, and limited MERS-CoV data implicate NSP8 as the main RNA binding protein in the RTC, driven by the positively charged NSP8 “sliding poles” which interact with the negatively charged RNA backbone. In keeping with this, upon addition of individual and pairwise combinations of NSPs, we found that of the individual NSPs, NSP 8 alone strongly produced a shift in the fluorescent RNA band, implying RNA binding (Figure 1E&F, lane 3). Addition of NSP7+8 and NSP8+12 also resulted in RNA binding, however at lower levels than that observed for NSP8 alone (Figure 1E&F, lane 5). In agreement with the denaturing assay, addition of the full RTC (NSP7+8+12) resulted in the highest level of RNA binding (Figure 1E&F, lane 8), suggesting that NSPs 7, 8, and 12 are all necessary for the formation of a transcriptionally active and stable RTC.

### The MERS-CoV minimal RTC extends RNA in a dose-dependent manner

After exploring binding and extension activity of the minimal MERS-CoV RTC through bulk assays, we employed an orthogonal single-molecule method to confirm fluorescent primer extension. Single-molecule Förster resonance energy transfer (FRET) measures the transfer of energy from a donor fluorophore to an acceptor fluorophore over a 2-10 nm range. The efficiency of this energy transfer (E) can be converted to a distance measurement based upon the fundamental properties of the donor-acceptor pair (29), providing a useful measurement of protein and/or RNA dynamics.

The same fluorescent template/primer duplex used in the in vitro experiments in Figure 1 was used for smFRET experiments as the template forms a self-hybridising overhang that can report on primer extension. In the non-extended duplex, Cy3 and Cy5 fluorophores, positioned after the 43^rd^ nucleotide of the template and between the 12^th^ and 13^th^ nucleotides of the primer respectively, were predicted to produce a high FRET efficiency due to their close proximity, while primer extension from addition of the minimal RTC and nucleotides would be expected to produce a lower FRET efficiency due to the template overhang opening (Figure 2A, (26)). As predicted, the annealed template-primer duplex alone produced a high apparent FRET efficiency of 0.87 (Figure 2A).

**Figure 2:**
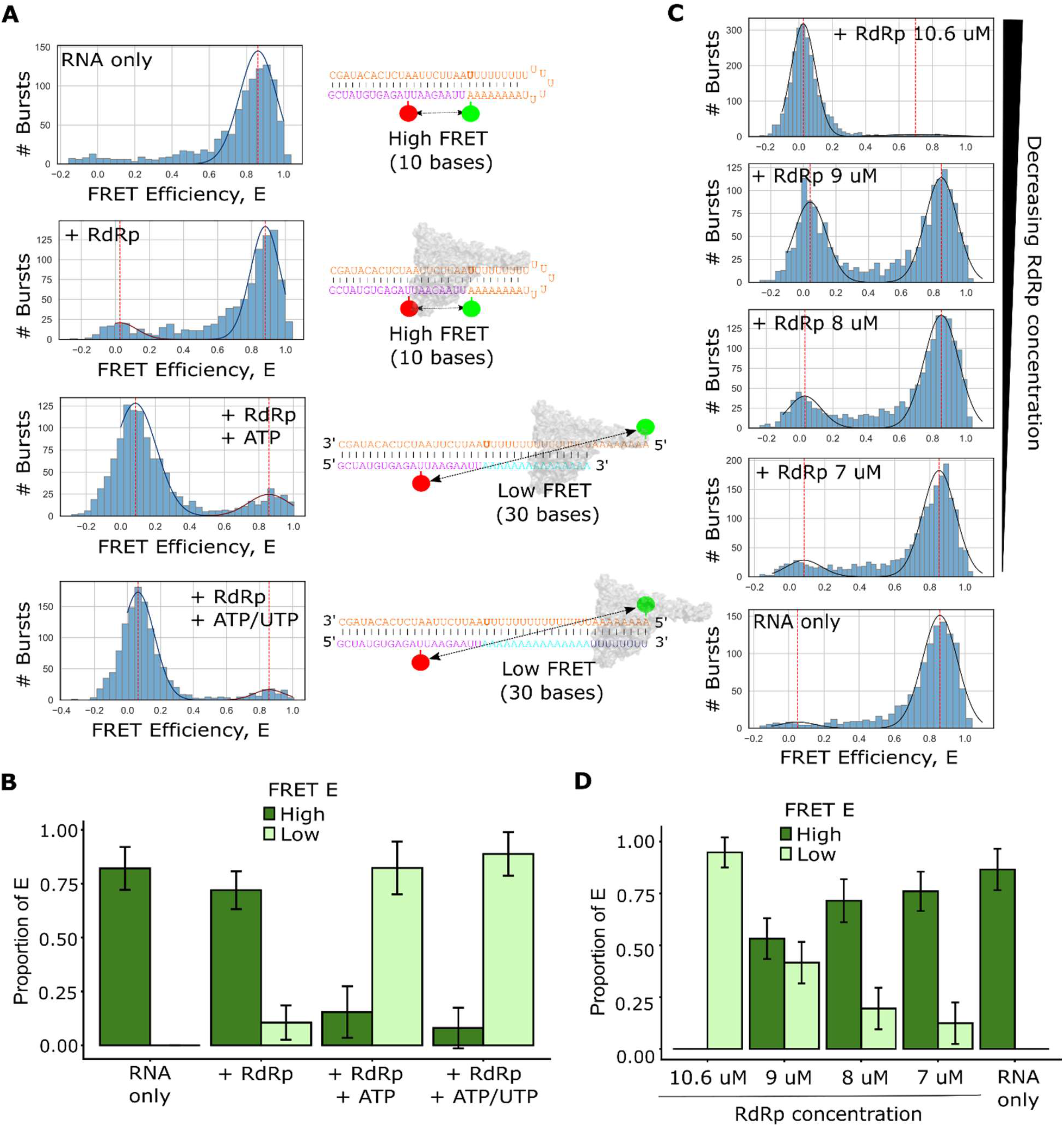
Characterisation of the MERS-CoV RTC using a smFRET reporter assay. **A)** FRET efficiency (E) histograms of nucleotide dependent extension of the MERS RdRp. The mean of each fitted population is represented by a red line. To the right of each histogram is a schematic representation of the RNA in each condition. Annealed RNA forms a self-annealing hairpin that brings the Cy3 and Cy5 dyes close in proximity and results in a high FRET population. Upon RdRp addition, no change is detected in transfer efficiency, whilst further addition of NTPs leads to an unfolding of the template due to extension, increasing the distance between the dyes and producing low FRET populations. **B)** Quantification of FRET efficiency histograms of nucleotide dependent extension as a proportion of the RNA population. Error bars correspond to the standard deviation of the gaussian distributions. High FRET populations are indicated in dark green and low FRET populations in light green. **C)** Decreasing concentrations of the MERS-CoV RdRp reduce the extended low FRET efficiency RNA population. **D)** Quantification of FRET efficiency histogram plots of RTC titration, as a proportion of extended RNA. Error bars correspond to standard deviation of the fitted gaussian distributions.

Upon incubation of the fluorescent RNA with the minimal MERS-CoV RTC, in the absence of NTPs, we again observed a high apparent FRET efficiency of 0.89, confirming nucleotide dependence. The addition of the minimal RTC and ATP produced a major shift in the population from high FRET (15%) to a low apparent FRET efficiency of 0.08 (85%), indicating that a large proportion of primer-template RNA duplexes were extended by the MERS-CoV minimal RTC (Figure 2A&B). Upon addition of UTP alongside ATP, the proportion of fully extended RNAs shifted to 92% (Figure 2A&B). We were thus able to confirm that the MERS-CoV RTC was capable of completely extending RNA templates in smFRET assay conditions.

To understand the effects of RTC concentration on the proportion of extended RNA, we titrated the RTC concentration down, keeping individual NSP ratios constant. Whilst the addition of 10.6 µM of RTC led to a near-total extension of the RNA population, reduction of RTC concentration to 9 µM led to a ∼2-fold reduction in extension (Fig 2C&D). Upon the addition of 7 µM RTC this was reduced to 20%, indicative of the extension of the template-primer RNA being dose-dependent relative to the MERS-CoV RTC concentration.

### Cryo-EM reveals two distinct RNA-bound MERS-CoV polymerase complexes

To obtain the structure of the MERS-CoV replication transcription complex we mixed recombinant NSP 12, 7, and 8 at a ratio of 1:2:4. RNA mimicking template and primer were annealed at a 1:1 ratio and subsequently added for a final molar ratio of 1:1 to NSP12. A single particle cryo-EM dataset was subsequently collected. The reconstructed maps showed to two distinct MERS-CoV complexes (Supplementary Figure 1). One complex containing one copy of NSP12, one copy of NSP8 (NSP8a) and both viral RNAs was determined to a resolution of 3.27 Å (Figure 3A&B). A second complex containing an additional copy of NSP8 (NSP8b) and a copy of NSP7 was determined to a slightly lower resolution of 3.41 Å (Figure 3A&C).

**Figure 3:**
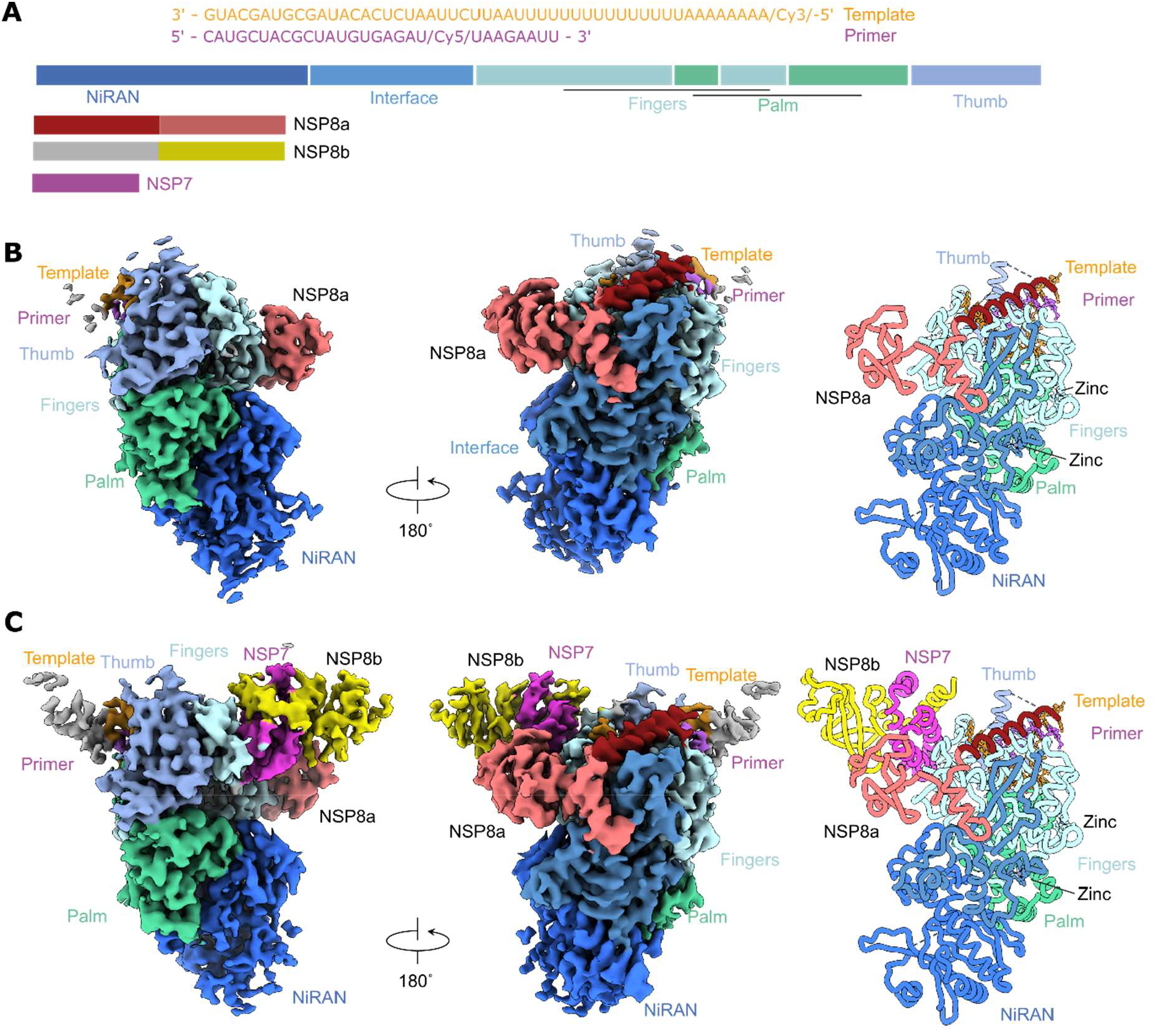
Cryo-EM structures of the MERS-CoV RTC. **A)** Domain architecture of MERS-CoV NSP12, 7 and 8 and RNA templates. **B)** Map and model of the NSP12/8a complex and both viral RNAs with domain annotations. **C)** Map and model of the NSP12/8a/8b/7 complex and both viral RNAs with domain annotations.

Analysis of the structures revealed an N-terminal NiRAN domain, followed by an interface domain connecting to the C-terminal RdRp domain. Neither structure we determined contained a fully resolved NiRAN, potentially caused by the absence of the nucleotide substrate for this domain. The RdRp domain within NSP12 contained the canonical RNA and NTP binding motifs A-F, with the two active site residues D763/D764 contained within motif C (Supplementary Figure 2A). Two non-catalytic ions, putatively identified as Zinc ions though coordination pattern and homology to other coronavirus NSP12 structures, were bound within the Interface and Fingers regions. Comparison of the two MERS-CoV RTC structures revealed notable differences in the organisation of the NSP12 active site. While the overall architecture of NSP12 was highly conserved between the partial NSP12/8a and complete NSP12/8a/8b/7 complexes, several regions surrounding the catalytic centre exhibited differences in local structural order. In the partial complex, residues 444-449 within the RdRp fingers subdomain, together with the tip of motif F, were disordered and could not be resolved in the cryo-EM density. In contrast, both regions were clearly resolved in the complete RTC, indicating increased structural ordering upon incorporation of NSP7 and NSP8b (Supplementary Figure 2B). Notably, motif F contributes to the formation of the NTP entry channel and contacts the RNA template within the polymerase active site (Supplementary Figure 2B).

Bound to the central NSP12 are two copies of NSP8, a single copy of NSP7, and regions of the template and primer RNAs (Figure 3A&C). NSP8a is bound directly to the Interface and Fingers domain of NSP12, with only the C-terminal region (residues 79-193) resolved in the model. The interaction of 2241 Å^2^ forms the largest interface between components, with this large interface likely the reason it is present in both complexes we determined. NSP7 forms the next major contact to NSP12 (burying 631 Å^2^) with the interface mediated through the Fingers region and a minor interface contacting NSP8a. NSP8b is maintained in the complex through interaction with a large surface area of 1586 Å^2^ to NSP7 and a much smaller 102 Å^2^ interface directly with NSP12. Only the C-terminal region of NSP8b is ordered in this complex (Figure 3A&C).

In addition to the protein subunits, we observe two RNA molecules - one acting as the template RNA and the other as the product or primer RNA (Supplementary Figure 2C). We can clearly resolve bases 21-32 of the template and bases 21-28 from the primer RNA. We observe a tight RNA duplex exiting the RdRp core and a 3-base overhang positioning U23 in the active site (Supplementary Figure 2C). We do not observe the extended template-product duplex observed for other coronavirus RTC structures, potentially because of the insertion of fluorescent molecules into the RNA backbone of both the template and primer, which could break the duplex. In the absence of the RNA duplex, the N-terminal region of both NSP8s, which forms an extended helix in other structures, is disordered.

### MERS-CoV NSP12 is compatible with SARS-CoV-2 NSP7 and NSP8, but not vice versa

Comparison between our newly solved MERS-CoV RTC structure and the SARS-CoV-2 RTC solved by Hillen et al. (17) confirmed that the two complexes share high structural similarity. Although the N-terminal region of the NSP8s is disordered in the MERS structure, there is an overall root mean square deviation (RMSD) of 0.84 Å between equivalent C-alpha carbons in the two structures, suggesting strong conservation (Figure 4A). Comparison of the MERS-CoV and SARS-CoV-2 RTCs reveals a broadly conserved area of inter-subunit interactions, although the sizes of individual interfaces differ. The NSP12–NSP8a and NSP7–NSP8b interfaces are larger in MERS-CoV than in SARS-CoV-2 (2,241 vs 1,868 Å^2^ and 1,586 vs 1,352 Å^2^, respectively), whereas the NSP12–NSP7 interface is smaller (631 vs 785 Å^2^). Thus, despite differences in the extent of buried surface area, the major interaction interfaces are maintained between the two RTCs.

**Figure 4:**
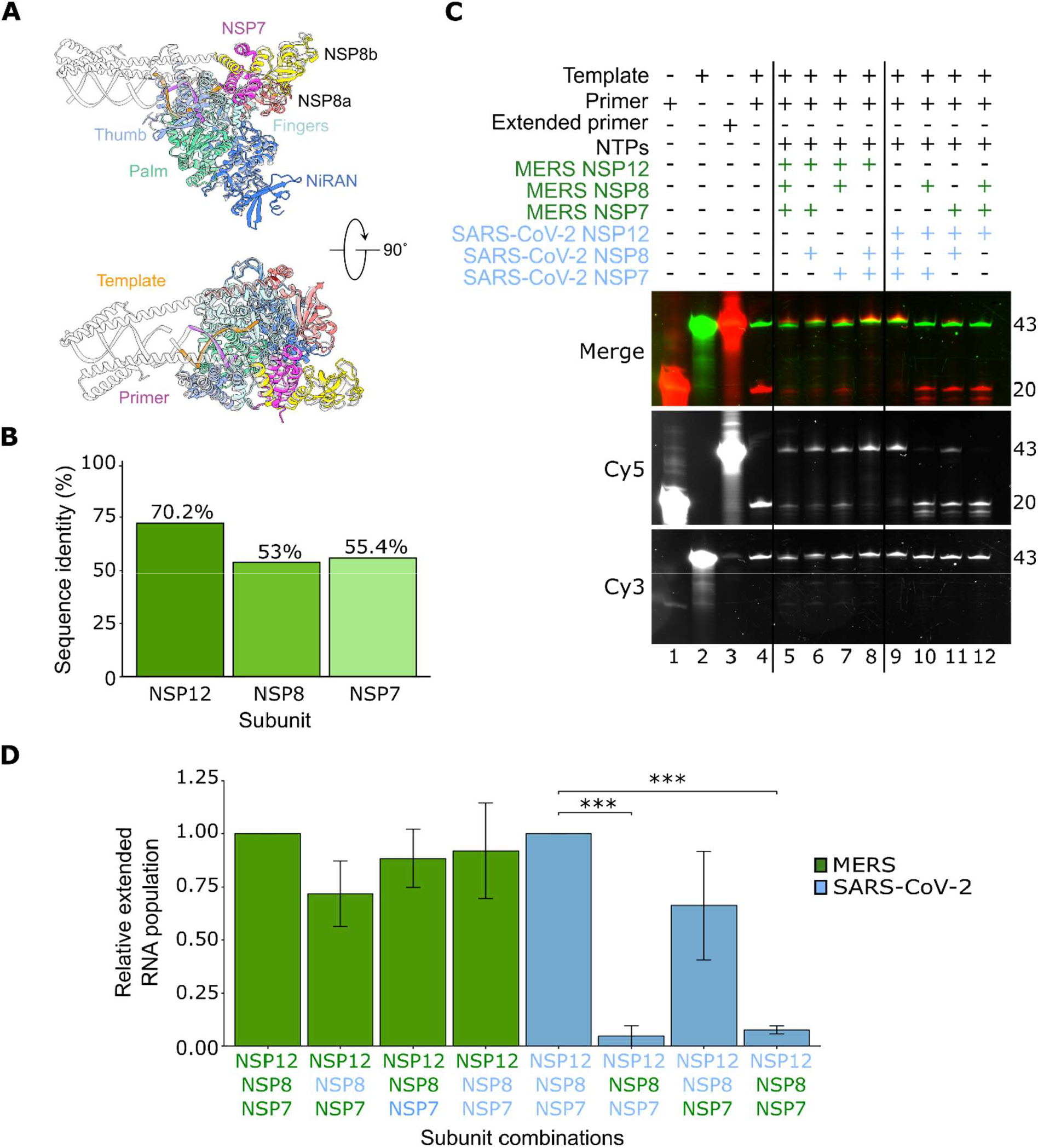
An in-vitro extension assay of MERS-CoV and SARS-CoV-2 subunit swapping. **A)** Comparison between the MERS-CoV RTC structure and the SARS-CoV-2 RTC (17) confirming that the two complexes share high structural similarity (RMSD of 0.84Å). MERS is in colour, whilst SARS-CoV-2 is in transparent grey. **B)** Amino acid sequence similarity between SARS-CoV-2 and MERS-CoV NSPs. **C)** Subunit swapping explored the compatibility of MERS and SARS-CoV-2 NSPs in complex formation and extension activity. **D)** Quantification of the subunit swapping extension assay. NSP combinations with the MERS NSP12 are shown in green, normalized to the control assay, while NSP combinations with the SC2 NSP12 are shown in blue and normalised to the control assay. Error bars represent the standard error of the mean of a minimum of 3 independent repeats. Statistical analysis was conducted using student’s unpaired t-test.

A pairwise sequence alignment of the MERS-CoV and SARS-CoV-2 NSPs used in the minimal RTC extension assays, revealed an amino acid sequence identity of 70.2%, 53% and 55.4% for NSPs 12, 8 and 7 respectively (Figure 4B, Supplementary Figures 3-5). This is relatively low compared to the previously published 98.8%, 97.4% and 96% identity between SARS-CoV and SARS-CoV-2 NSPs 12, 8 and 7 (30). Previous efforts to understand the extent of this sequence variation through SARS-CoV and SARS-CoV-2 subunit cross-combination assays showed that NSP8 replacement significantly affected NSP12 mediated activity, whilst NSP7 had a minimal effect (31). Considering the comparable structures of the minimal RTCs and inhibition of both the SARS-CoV-2 and MERS-CoV RTCs by nucleoside analogue Remdesivir (22), (32), we questioned whether the MERS-CoV and SARS-CoV-2 RTCs would also support NSP cross-combination.

We combined analogous MERS-CoV and SARS-CoV-2 NSPs in an in vitro extension assay, producing combinations of the three essential NSPs at or above the stoichiometric 1:5:5 ratios reported earlier (Figure 4C). Upon addition of either SARS-CoV-2 NSP 7 or 8 with a MERS-CoV NSP 12 (Figure 4C lanes 6-8, 4D), we observed that the addition of SARS-CoV-2 NSP7 produced a comparable level (88 +/-13.6%) of extension activity to the native MERS-CoV NSPs. However, SARS-CoV-2 NSP8 replacement reduced the activity of the RTC slightly to 72 +/-15.5% of the native MERS-CoV RTC. Addition of both SARS-CoV-2 NSPs to MERS-CoV NSP12 resulted in activity that was roughly comparable (92 +/-22.5%) to the native MERS-CoV RTC.

In contrast, assays with MERS-CoV NSP7 addition to the SARS-CoV-2 RTC produced a stronger reduction in activity than addition of SARS-CoV-2 NSP7 to the MERS-CoV RTC (66% +/-25.5%) (Figure 4C lane 11, 4D). The largest reduction in activity came from the addition of MERS-CoV NSP8 to the SARS-CoV-2 RTC, where extension activity was reduced to just 5% +/-4.8% of that observed during SARS-CoV-2 RTC extension. Addition of both MERS-CoV NSP7 and NSP8 to SARS-CoV-2 NSP12 resulted in extension activity at only 8% +/-1.89% of wildtype activity. Whilst substitution of native NSP7 therefore reduced activity partially across both systems, native NSP8 substitution had a significant effect; this is not surprising considering the role that NSP8 forms in binding genomic RNA and interfacing with NSP12 prior to initiation. This reveals asymmetric compatibility and identifies NSP8/NSP12 interactions as an area of virus-specific adaptation despite overall RTC conservation.

Next, we sought to rationalise the molecular basis for MERS-CoV NSP8 being unable to support SARS-CoV-2 NSP12 activity. As addition of MERS-CoV NSP7 restored 66% of activity, but addition of both MERS-CoV accessory proteins did not, we reasoned that the NSP12-NSP8a interface was likely not being appropriately reconstituted in the mixed complex. Analysis of the interface regions from each complex revealed two potential regions causing this mismatch (Supplementary Figure 6). SARS-CoV-2 NSP12 K391 is a leucine at the equivalent 394 position. The pocket where this residue is positioned is more hydrophobic (T137/T141 to V138/V142) in the MERS-CoV NSP8, providing a less favourable environment for binding of K391. In addition to these, the cumulative effect of several smaller changes (Supplementary Figure 7) may weaken the interface such that the chimeric complex is less stable.

### Suramin, a broad-spectrum inhibitor of viral RdRps, inhibits MERS-CoV RTC at low micromolar concentrations

Our cryo-EM structure and paired sequence alignment showed that the core catalytic regions of the MERS-CoV RTC complex are highly conserved between SARS-CoV-2 and MERS-CoV. We therefore hypothesised that known inhibitors of the SARS-CoV-2 RTC might also work on MERS-CoV. Suramin, originally proposed as an antiparasitic drug, has been shown to provide broad-spectrum inhibition of viral infection at cell entry and viral particle release stages (33), (34). Recent biochemical data has shown that suramin is a potent inhibitor of the SARS-CoV-2 RTC (26). Within the RdRp structure, two suramin molecules bind active site motifs that usually make contacts with the RNA template-primer duplex, thus blocking binding of the RNA and preventing entry of NTPs into the active site (19).

We titrated increasing concentrations (2.34 - 300 µM) of suramin into our in vitro denaturing extension assay alongside the RTC, NTPs and the template-primer duplex. Upon addition of suramin, the 43 nt extension product reduced in a dose-dependent manner, with visible extension product signal being lost entirely between the 19 µM and 38 µM suramin concentrations (Figure 5A, lanes 8-9). We then confirmed this reduction using our smFRET assay, introducing suramin alongside RNA and NTPs. We reduced our suramin titration to a range of 0.75 – 100 µM (Figure 5B&C). Low FRET efficiency extension products were less than 50% of the burst population upon addition of 0.75 µM suramin, and at a concentration of 100 µM suramin no low FRET population was observed (Figure 5B&C). We therefore confirmed that suramin can inhibit the MERS-CoV RTC in a concentration dependent manner.

**Figure 5:**
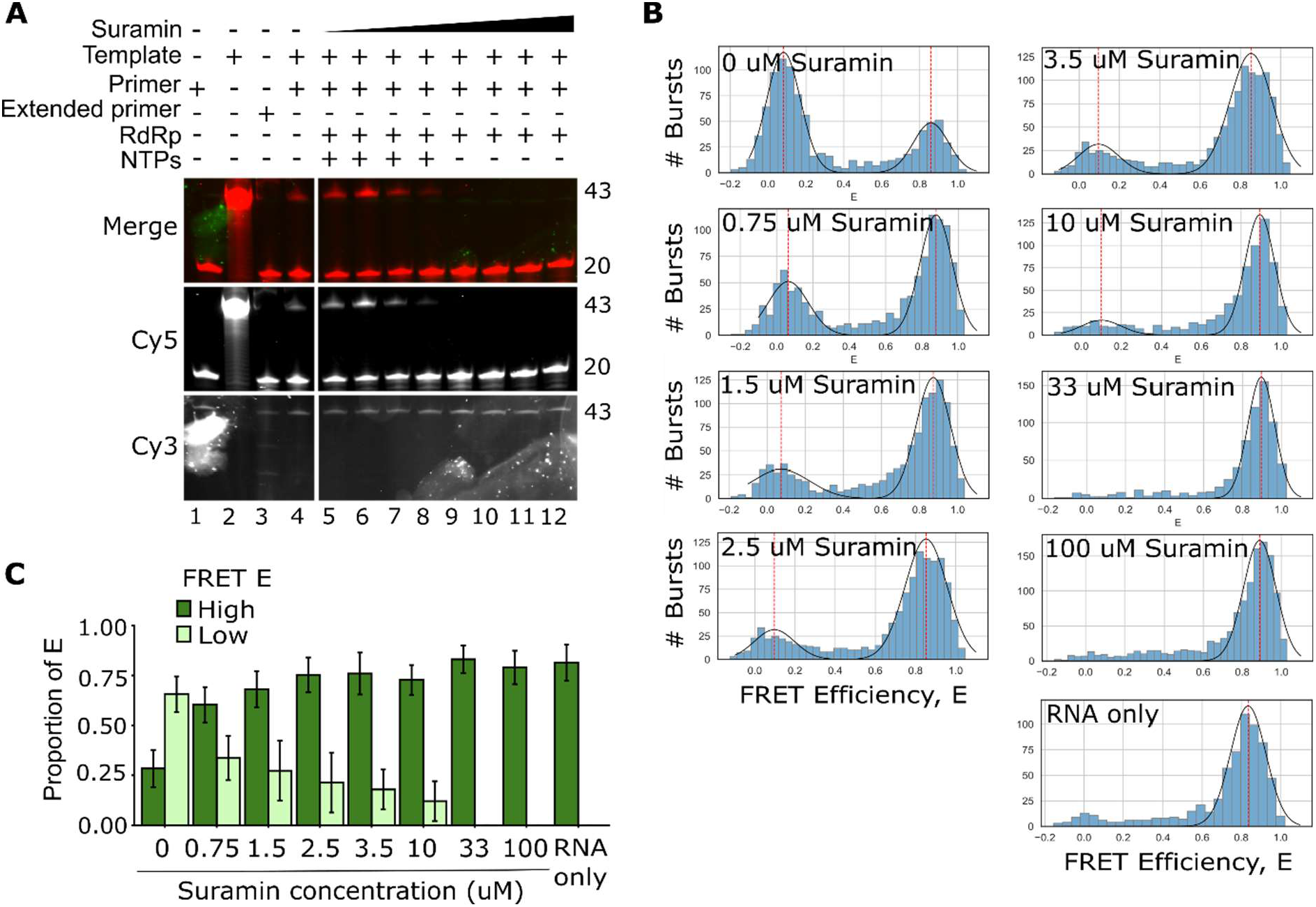
Inhibitory effects of Suramin on MERS RTC extension activity *in-vitro* and at the single molecule level. **A)** A representative *in-vitro* titration of Suramin in the presence of a fixed concentration of MERS RTC and NTPs. Control lane 4 contained RNA, RTC (9.6 µM) and NTPs (500 µM). Suramin was titrated from 2.34 µM to 300 µM (Lanes 5-12). We note a spill of extended primer control (lane 2) present in lanes 3 and 4, unrelated to extension activity. **B)** FRET efficiency (E) histograms of a Suramin titration assay using smFRET. Suramin was introduced at 0.75 µM, up to 100 uM, with the RTC concentration at 8.58 uM and NTPs at 500 uM. A bimodal distribution of non-extended and extended RNA populations is accentuated by the presence of Suramin, with high concentrations completely abolishing low FRET E populations. Gaussian distributions were fitted to RNA populations with mean represented by red line. **C)** Quantification of FRET efficiency histograms in 5B, as a proportion of extended RNA. Error bars correspond to the standard deviation of the fitted gaussian distributions.

## Discussion

While SARS-CoV-1 and SARS-CoV-2 have been extensively studied following their respective epidemic and pandemic outbreaks, MERS-CoV, which emerged in the intervening period, remains comparatively poorly characterised at the biochemical level, particularly with respect to its core replication machinery. Towards filling this gap, our fluorescent in vitro assays, combined with smFRET data, have confirmed several key findings regarding the formation and composition of the MERS-CoV replication-transcription complex. First, we have been able to confirm that the minimal RTC is formed of NSPs 7, 8, and 12, consistent with the limited preliminary data available. (18). We also note that NSP8 and 12 together can form a complex capable of templated primer extension, albeit at heavily reduced extension activity than the minimal RTC, in partial agreement with the findings of Gordon et al. (32).

Due to the lack of partial extension products in our NSP8/12 assay (Figure 1C, lane 9), the MERS-CoV NSP7 forms a key regulatory step in transitioning the RTC either from an unstably bound complex likely to dissociate from the RNA prior to initiation to a stable complex, or from a stably bound complex incapable of initiating the first polymerisation step consistently, to one capable of initiating primer extension. Recent single-molecule protein-induced fluorescence enhancement (smPIFE) assays have suggested that the SARS-CoV-2 NSP7 does not appear to improve complex stability, but acts as an inducer of polymerisation, perhaps akin to the bacterial RdRp sigma factor (27), (35). Single-molecule magnetic tweezer data further supported this, suggesting that it was a conformational change rather than RNA binding or RTC assembly that delayed polymerase activation (36). It is therefore possible that this characteristic of NSP7 is consistent across coronavirus RdRps.

We also confirmed that NSP8 acts as the key RNA binding component of the MERS-CoV RTC, similarly to SARS-CoV-2 and SARS-CoV RTCs. We did not detect binding of NSP12 or 7 in the absence of NSP8. It is likely that a significant proportion of NSP8 binding is non-specific – the NSP8 structure does not appear to recognise specific RNA sequences in the absence of NSP12, as the negatively charged N-terminal domain alone only can bind duplexed RNA strongly (37). Although the polyprotein cleavage step of initial viral translation upon entry appears to suggest a pre-assembly of the RTC through interactions of NSP 8, 7 and 12, it is also possible that coronavirus replication-transcription complex formation is a stepwise process, driven by the binding of NSP8 to dsRNA of a minimal length, leading to recruitment of NSP12, and then subsequent transition into initiation either through a change in conformation, or interactions with NSP7 which drives conformational change and the first polymerisation step. NSP7 could then either remain with the polymerase or dissociate near-instantly.

Our cryo-EM structures provide new insights into the molecular architecture and assembly of the MERS-CoV replication-transcription complex, revealing a high degree of structural conservation with the SARS-CoV-2 RTC despite substantial sequence divergence between the two viruses. Structural alignment with the SARS-CoV-2 RTC described by Hillen et al. (17) revealed an overall RMSD of 0.84 Å, demonstrating that the organisation of the polymerase core and its accessory subunits is strongly conserved. Interestingly, we resolved two distinct RNA-bound complexes: a partial complex containing NSP12 and a single copy of NSP8 (NSP8a), and a complete complex containing NSP12, NSP7, and two copies of NSP8. The presence of NSP12–NSP8a in both structures, together with its extensive buried interface, suggests that this interaction forms a particularly stable component of the RTC and may represent an early intermediate in complex assembly. In contrast, NSP7 and the second copy of NSP8 (NSP8b) were observed only in the complete complex, where NSP8b interacts predominantly with NSP7 through a substantial interface, while making relatively limited direct contact with NSP12. These observations are consistent with a potential stepwise assembly mechanism in which NSP12 first associates with NSP8a, followed by recruitment of an NSP7– NSP8b heterodimer, although the spatial and temporal order of these interactions remains to be established.

By combining SARS-CoV-2 and MERS-CoV proteins into a single assay, we have shown that some of these interactions appear to be virus-specific. First, using a denaturing extension assay, we established that the MERS-CoV NSP12 can utilize SARS-CoV-2 NSPs with a negligible loss of activity, whilst SARS-CoV-2 NSP12 struggles to utilize MERS-CoV NSP8. This effect was not rescued by simultaneous substitution of MERS-CoV NSP7, suggesting that incompatibility between heterologous NSP7 and NSP8 alone is unlikely to explain the reduction. Instead, our cryo-EM structure identifies the NSP12–NSP8a interface as the largest intersubunit contact involving NSP12, suggesting that differences in this interface may contribute to the observed asymmetry. Previous studies have demonstrated that substitutions affecting hydrophobic and polar interactions at the SARS-CoV-2 NSP12–NSP8 interface can substantially impair polymerisation activity (38). Sequence differences between MERS-CoV and SARS-CoV-2 NSP8 may therefore alter the stability or geometry of this interface, with MERS-CoV NSP12 potentially displaying greater tolerance to heterologous NSP8 binding. Alternatively, the loss of activity may reflect impaired conformational activation rather than an inability to assemble the complex. Further structural and kinetic characterisation of hybrid RTCs will be required to distinguish impaired complex assembly from defective polymerase activation.

The increased ordering of this region in the complete complex suggests that recruitment of NSP7 and NSP8b may stabilise the active-site architecture, promoting the correct positioning of the RNA template and incoming nucleotides for efficient RNA synthesis. This observation is consistent with our biochemical findings, in which the NSP12–NSP8 complex exhibited limited primer extension activity (∼8%), whereas addition of NSP7 substantially increased polymerisation activity. Together, these findings suggest that NSP7 and NSP8b may contribute to polymerase activation by stabilising key structural elements surrounding the catalytic centre, rather than functioning solely as accessory scaffolding proteins. However, whether this ordering directly results from cofactor binding or reflects differences in conformational states captured during cryo-EM reconstruction remains to be established.

Finally, we utilized our smFRET assay to characterise the extension activity of the MERS-CoV minimal RTC. As has been previously discussed, the overhang duplex smFRET assay enables a rapid method of studying antiviral inhibitors, and as predicted in the Groves et al., (26) article, this overhang hairpin assay can screen multiple coronavirus RdRps in addition to the SARS-CoV-2 RdRp. We have further confirmed that MERS-CoV, like SARS-CoV and SARS-CoV-2, is capable of inhibition by the non-nucleoside inhibitor suramin, further opening the door to utilization of novel suramin mimics against the MERS-CoV virus. The structural conservation observed between the MERS-CoV and SARS-CoV-2 RTCs, together with their partial functional interchangeability and shared susceptibility to polymerase inhibition, highlights the RTC as a promising target for broad-spectrum antiviral development.

Together, our findings address a substantial gap in the understanding of MERS-CoV replication by providing a structural and biochemical framework for its core replication machinery. By defining the composition and functional roles of the minimal RTC, determining its RNA-bound structure, and identifying both conserved and virus-specific features of polymerase–cofactor interactions, this work extends our understanding of coronavirus replication beyond the extensively studied SARS-CoV-2 system. Importantly, the MERS-CoV RTC structure provides a molecular template for structure-guided inhibitor discovery and optimisation, while the biochemical and single-molecule assays established here provide complementary platforms for evaluating candidate compounds. Given the continuing circulation and pandemic potential of MERS-CoV, and the capacity of coronaviruses to evolve through mutation and recombination, expanding our molecular understanding beyond currently dominant human coronaviruses is essential for pandemic preparedness. This work therefore provides both fundamental insight into MERS-CoV RNA synthesis and an experimental foundation for the development of antiviral strategies against MERS-CoV and future emerging coronaviruses.

## Materials and Methods

### RNA

All RNA constructs were ordered from Integrated DNA Technologies (IDT) based on sequences utilised previously in Groves et al., (26) conjugated to fluorescent dyes Cy3 and Cy5 through phosphodiester bonds with bases (Supplementary Table 1). Subsequent rounds of nucleotide addition occurred to produce the full-length constructs after fluorophore conjugation, dependent on internal or end-labelled positioning. Template and primer RNAs were annealed at a final concentration of 300 nM in an annealing buffer consisting of 20 mM Tris-HCl pH 8.0, 1 mM ethylenediaminetetraacetic acid (EDTA), 500 mM NaCl. The annealing step used a thermocycler to reduce temperature from 95°C to 4°C in a series of steps, following a single cycle. The temperature was reduced at 2.5°C steps from 95°C to 65°C for two minutes per step, followed by a 5-minute step at 65°C, a 10-minute step at 45°C, a 5-minute step at 25°C, and held at 4°C thereafter.

### Protein constructs

The NSP12 ORF in a pFastBac DualFplasmid was purchased (Synbio Technologies) before being transformed into DH10 MultibacCre cells. Successful colonies were identified using blue-white screening. Recombinant bacmids were then isolated from the DH10 MultibacCre cells before being transfected into Sf9 cells, to produce a V0 and subsequent V1 baculovirus stock. Subsequent V1 baculovirus was used to infect Sf9 cells. These were incubated shaking at 27° C at 120 rpm for 3 days, harvested by centrifugation for 15 mins at 2500 g at 40°C, to yield pellets expressing NSP12 which were stored at -80° C.

Initial plasmid constructs carrying GFP-tagged NSP7 and NSP8 genes in a pCAGGS background were provided by Dr Christopher Coleman, University of Nottingham, who obtained them from Dr Matthew Frieman at the University of Maryland School of Medicine, where they were originally cloned. To produce plasmids for E. coli expression, the plasmid pGEX-6P-1 backbone was utilized. This backbone was linearised by a double restriction digest using BamH1 and Xba1 (NEB). PCR and Gibson Assembly were used to assemble the NSP genes into pGEX-6P-1. Sequence verified plasmidswere transformed into E. coli RIPL cells, plated onto LB plates with 35 µg/mL chloramphenicol, 50 µg/mL streptomycin, and 100 µg/mL ampicillin, before upscaling growth in volumetric flasks. When OD had reached ∼0.4-0.6, 1 mM IPTG was added to induce expression of proteins and E. coli harvested after 16 hours growth at 18°C.

### Protein purification

Proteins were purified as in our previous study. One litre of NSP12 containing sf9 cells was resuspeneded in wash buffer [50 mM HEPES-NaOH, pH 7.5, 300 mM NaCl, 10% glycerol, 0.05% n-Octyl β-d-thioglucopyranoside (OTG), 1 mM dithiothreitol (DTT), 1 protease inhibitor tablet] and lysed by sonication (26). Cell lysate was clarified by centrifugation at 35,000 x g at 4°C for 30 minutes. The resulting supernatant was combined with IgG sepharaose resin and incubated for 3 hours at 4°C. Resin was then washed with 20 volumes of wash buffer before addition of TEV protease for 16 hours at 4°C. The eluate was concentrated and applied to a superdex200 increase 10/300 GL column equilibrated in 25 mM HEPES, pH 7.4, 100 mM NaCl, 2 mM MgCl2,1 mM DTT.

One litre of NSP7 or NSP8 was resuspended in wash buffer further supplemented with 50mg lysozyme and 100 µg/mL RNAase A prior to lysis by sonication. Supernatant was clarified, applied to GST Sepharose, and incubated for 3 hours at 4°C. Resin was washed with 20 column volumes of wash buffer prior to addition of 3C protease overnight at 4°C. Eluted protein was then applied to a superdex75 increase 10/300 GL column equilibrated in 25 mM HEPES, pH 7.4, 100 mM NaCl, 2 mM MgCl2,1 mM DTT.

All proteins were assessed for purity by SDS-PAGE and frozen in liquid nitrogen before storage at -80°C.

### Electrophoretic mobility shift assays

Electrophoretic mobility shift assays (EMSAs) were performed using annealed RNA at a 50 nM final concentration. RNA was mixed with a master mix containing 5 mM MnCl_2_, 0.5 mM NTPs, 10 mM KCl, 1U/μl RNasin (Promega), 0.1 mM DTT, a pre-incubated mixture of NSPs 12, 7 and 8 at a ratio of 1:5:5 or greater, and Protein Storage Buffer (PSB, 6.25 mM HEPES, 37.5 mM NaCl, 0.5 mM MgCl_2_, 0.25 mM DTT, 20% glycerol). Pre-incubated NSPs were left on ice for 15 minutes prior to addition to the reaction mixtures. Reaction mixtures were then incubated at 30°C for 30 minutes prior to mixing 1:1 with native loading buffer (62.5 mM Tris-HCl pH 6.8, 40% Glycerol) and then loaded onto a 4-15% gradient Mini-PROTEAN TGX gel (Bio-Rad) in Tris-Glycine running buffer (25 mM Tris pH 8.8, 192 mM Glycine). Samples were run at 100V for 60-80 minutes before visualization on Bio-Rad ChemiDoc using Cy5 and Cy3 excitation/emission ranges.

### In vitro extension assays

NSPs were preincubated on ice for a minimum of 15 minutes to achieve an approximate stoichiometric ratio of 1:5:5 (NSP 12:8:7). A reaction mix consisting of PSB, master mix (5 mM MnCl_2_, 0.5 mM ATP/UTP, 10 mM KCl, 1 U/µl RNasin, 0.1 mM DTT), annealed RNA and preincubated NSPs was incubated for 30 minutes at 30 °C. Samples were then combined 1:1 with formamide and boiled for 3 minutes at 95 °C. Samples were loaded onto a 20% 19:1 Acrylamide/Bis acrylamide 6M urea gel in 1x TBE buffer. Gel electrophoresis was performed for 3.5 hours at 450 V before imaging in separate fluorescence channels, detecting fluorescent signal from Cy3 and Cy5 fluorophores. For the suramin (Sigma Aldritch) inhibition assay, the inhibitor was first diluted into water before subsequent dilution into PSB at 6-fold higher concentrations than reported. A final dilution step by addition to the reaction mixture produced concentrations stated in figures.

To quantify RNA binding or extension, we measured intensities of shifted and unshifted bands per lane using the Fiji gel measurement plugin (39). The intensities of bands corresponding to extended RNA (matching the length of control “extended” primer RNAs) and unextended RNA (matching the control primer RNA length) for extension assays, or bound and unbound RNA (matching the length of the unbound RNA control or shifted from this line) were compared to produce a ratio of bound/unbound or extended/unextended RNA. Graphs and statistical tests were plotted and performed in R.

### smFRET extension assays

To investigate the conformational states of polymerases during replication, SARS-CoV-2 overhang template RNAs with short primers were utilised to produce RNA duplexes. Our setup was described previously in Groves et. al., (26). Briefly, a 2-laser line was used to excite the donor and acceptor dyes in an EI-FLEX confocal microscope (Exciting Instruments). For donor excitation, a 520 nm laser, at 0.22 mW was used. Acceptor excitation utilised a 638 nm laser at 0.16 mW. Using Alternating-laser excitation (ALEX) over a 100 µs excitation period, the 520 nm laser followed a OFF:0 µs, ON:45 µs, OFF:55 µs frequency while the 638 nm laser followed a frequency of OFF:50 µs, ON:45 µs, OFF:5 µs. Exciting Instruments software was used for data collection and analysis.

Prior to the smFRET assays, NSP12,8 and 7 subunits were combined as with in vitro extension assays above. A final reaction mix, consisting of the NSP mix, master mix, PSB and 50 nM RNA, was incubated for 30 minutes at 30°C. Inhibitor assays incorporated compounds through dilution into PSB before addition to the reaction mixture. After incubation, reaction mixtures were diluted to ∼50 pM in confocal imaging buffer (5 mM MgCl_2_, 10 mM KCl, 0.5 U/µl RNAsin, 0.1 mM DTT, 25 mM HEPES, 100 mM NaCl and DEPC water) and run for 30 minutes.

### smFRET data analysis

Once captured by Exciting Instruments data collection software, datasets were processed using Jupyter notebook scripts, written by Exciting Instruments, utilising the open-source FRETBursts python package ((40), FRETBursts version 0.8).

Dataset correction factors α, δ, β, and γ were determined using “Correction Factor Finder Alpha-Delta” and “Correction Factor Finder Gamma-Beta” notebooks. These correction factors account for donor leakage into the acceptor channel, direct excitation of the acceptor fluorophore by the donor excitation laser, and system-based excitation and detection inequalities (41). After associating individual photons with the excitation periods listed above, a background rate was determined for an assay. An all-photon burst search with a minimum 6-fold higher-than-background threshold, followed by a dual-channel burst search, was applied to detected bursts to discriminate between background fluorescence and labelled molecules passing through the excitation volume. Stoichiometries outside of a 1:1 donor-to-acceptor ratio (0.2 < S <0.8) were eliminated to focus on dual-labelled molecules, producing one-dimensional FRET efficiency distributions.

We fitted the one-dimensional distributions with gaussian functions to produce mean FRET efficiencies for subpopulations of the distribution. Histograms were plotted as probability density functions, allowing us to determine the proportion of a given subpopulation relative to the whole population.

### Cryo-EM sample preparation and data collection

A complex of MERS-CoV NSPs 12, 7 and 8 bound to an annealed -28 RNA template and -28 primer duplex (Supplementary Table 1) was prepared at a 1:4.5:2:1 ratio, where the final complex concentration was 10.5 μM. NSPs were combined simultaneously, then RNA was added and incubated at room temperature for 1 hour. Grids were prepared using a Vitrobot mark IV (FEI) at 100% humidity, 10 blot force, and blotted for 3.5 seconds. Quantifoil R2/1 200 mesh grids were glow discharged, before a volume of 3.5 µL was applied and vitrified in liquid ethane/propane.

Cryo-EM data was collected at the Oxford Particle Imaging Centre on a 300 kV Titan Krios (Thermofisher Scientific) equipped with a SelectrisX energy filter and a Falcon 4 IV direct electron detector. Data were collected automatically using EPU V3.15, with a pixel size of 0.932 Å^2^ and a total dose of 47.9 e-/Å^2.^ in tiff format. 3000 movies were collected untitled before 7945 movies were collected with a 30° tilt.

### Cryo-EM data processing, structure determination, and model refinement

Data were processed using cryoSPARC V5.06 (42). Patch motion correction and patch CTF-estimation were performed with default setting. Movies with poor statistics were removed. An initial round of blob picking was used to generate picks for 2D classification. These templates were then used for template picking. The two particles sets were merged, duplicates removed, followed by 2D classification, and 3D reconstruction. Particles belonging to high resolution classes were then used to train a model in Topaz and a new particle set was picked (43). Particle sets from all approaches were then merged and again sorted using 3D classification and heterogenous refinement. Particles in high resolution classes were used to retrain topaz models twice more. A consensus reconstruction was then generated using non-uniform refinement, reference-based motion correction, and followed by local refinement. To identify the sub-populations containing nsp12/8A/RNA and nsp12/8AB/7/RNA iterative cycles of heterogenous refinement were used comprising maps of each complex and a junk class. Once particle populations were stable, a final round of local refinement was performed. A graphical scheme for processing is presented showing local map resolutions, particle distributions, and FSC (Supplementary Figure 1).

An initial model was determined using the AF3 server (44). The model was then manually positioned in the map in UCSF ChimeraX V1.12 (45) and large missing regions removed. Cycles of manual refinement in Coot (46) and real-space refinement in Phenix (47) were performed. Map and model statistics are described in Supplementary Table 2.

### Sequence alignment

EMBOSS needle (48) was used for the sequence alignment of RTC NSPs of MERS and SARS-CoV-2. Sequences for MERS-CoV subunits (England 1 strain) were used alongside the SARS-CoV-2 sequences from the Hillen et al. (17), 6YYT PDB structure. Protein sequences were aligned using the Needleman-Wunsch algorithm and BLOSUM62 matrix with the default gap open set to 10 and gap extend set to 0.5. Output from EMBOSS needle produced sequence alignments indicating full conservation, high conservation and non-conserved residues. Sequence identities reported therefore relate to full conservation of amino acids.

### Data, Materials, and Software Availability

Density maps and atomic models are available through the EMDB and the PDB with the following accession codes for structures determined in the study: MERS-CoV NSP12/8/RNA Complex (PDB 35GO / EMD-100513) and MERS-CoV NSP12/8/7/RNA Complex (PDB 35HQ / EMD-100533). Comparisons were made using a previously determined structure of a replicating SARS-CoV-2 polymerase (PDB 6YYT).

## Acknowledgements and funding sources

Author contributions: R.C., J.R.K., and N.C.R. conceptualised the study. R.C., J.R.K., N.C.R and S.S.P. wrote the manuscript. A.M.S., D.G., S.S.P., R.C., J.R.K., and A.D. performed experiments. S.S.P., J.R.K., and R.C. analysed and visualized the data. C.M.C., N.C.R. and J.R.K. contributed with project administration, supervision, resources, and funding acquisition. All authors commented on the manuscript and approved the submitted version. The work was funded by a Royal Society Dorothy Hodgkin Research Fellowship (DKR00620) to N.C.R., a BBSRC project grant (UKRI3595) to J.R.K and N.C.R., and an internal award from the School of Life Sciences at the University of Nottingham to C.M.C.

