## Supplementary material for "Molecular and Structural Characterisation of the MERS-Coronavirus Polymerase"

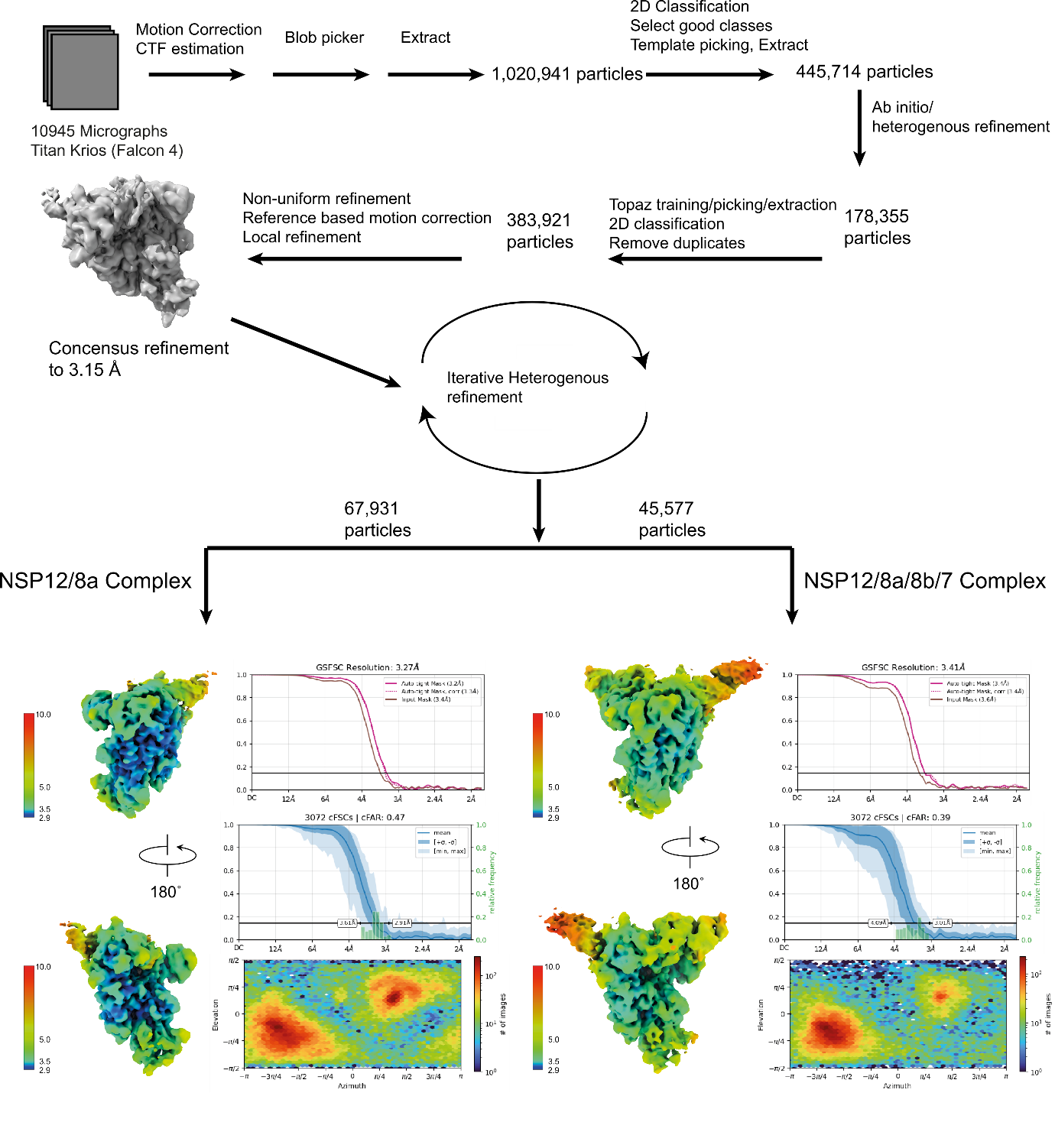

**Supplementary Figure 1.** **Cryo-EM processing scheme**. Scheme with final maps showing resolution estimations, local map resolutions, and particle distribution.

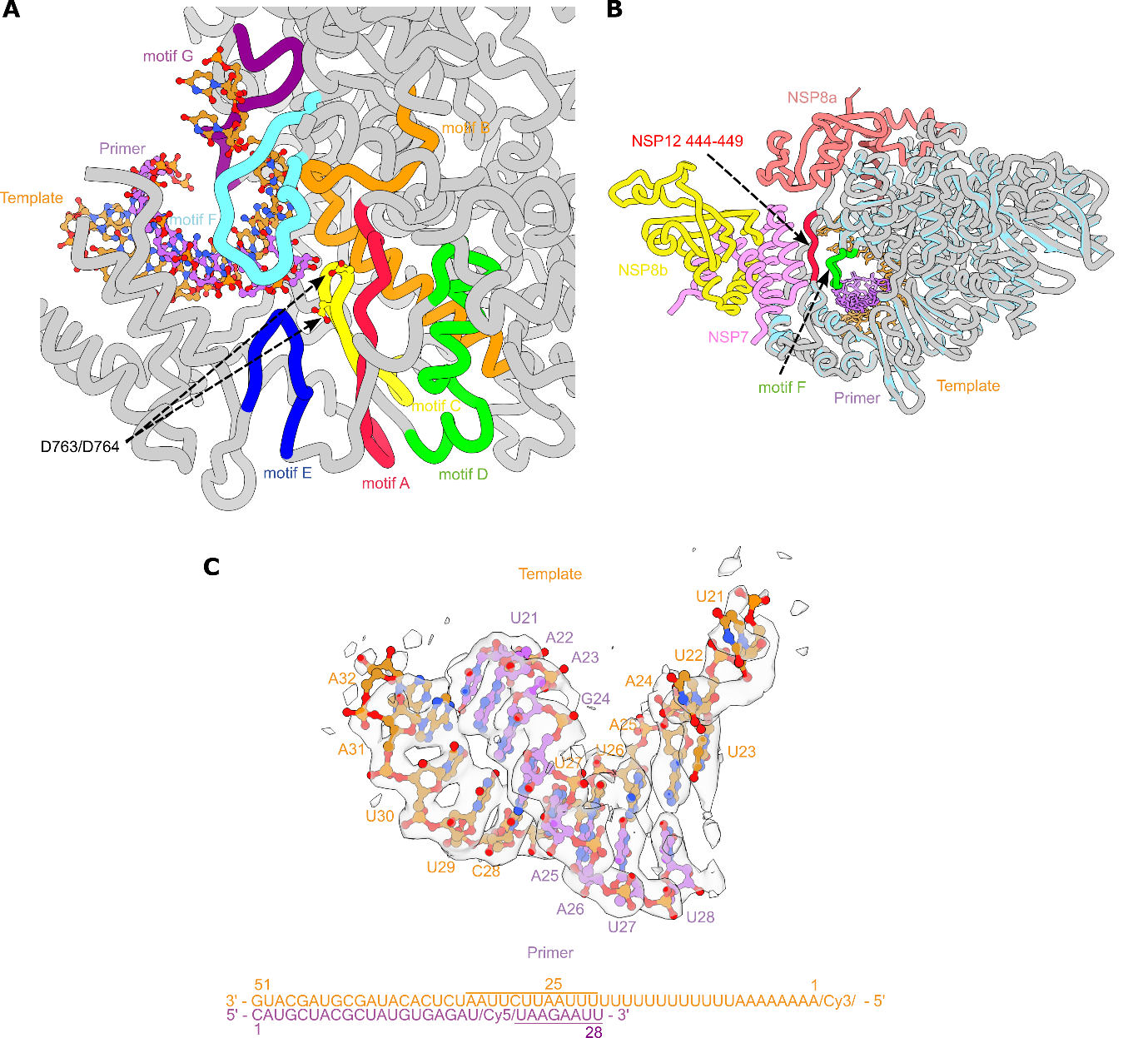

**Supplementary Figure 2.** **MERS-CoV RdRp active site analysis.** **A)** MERS-CoV NSP12 RdRp active site with RNA binding motifs annotated. NSP7, NSP8B, and residues 409-450 of NSP12 have been removed for clarity. Model shown is 35HQ/pdb_000035hq. **B)** Differences in the ordering of the active site when comparing the two structures. Residues 444-449 from the RdRp fingers (Red) and the tip of motif F (cyan) are disordered within the NSP12/8A structure. These regions are clearly visible within the NSP12/8A/9B/7 structure, where motif F forms part of the NTP entry channel and clamps the RNA template in the active site. The NSP12/8A complex is coloured light blue/cyan and the NSP12/8A/8B/7 complex is in grey (where not coloured by the other colours). **C)** Map density for each RNA segment with base number annotated.

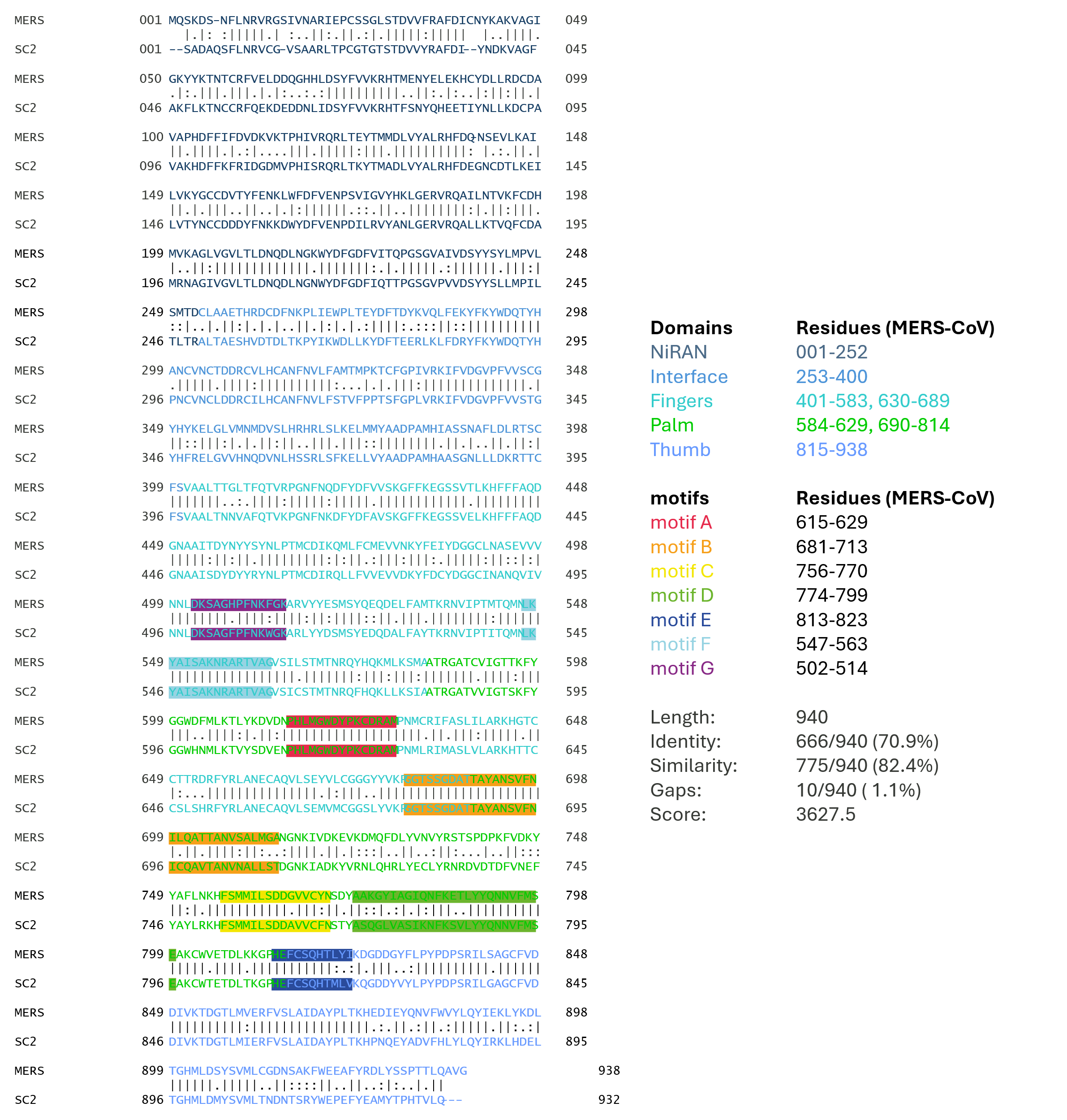

**Supplementary Figure 3.** **Sequence alignment of MERS-CoV and SARS-CoV-2 NSP12 using EMBOSS needle.** Highlighted residues indicate the SARS-CoV-2 motif boundaries (A-G) mapped onto MERS-CoV residues and coloured letters represent the domains: NiRAN (Dark blue), interface (blue), finger (teal), palm (green) and thumb (lighter blue) domains. Vertical lines depict identical residues, colons depict semi-conserved residues, and dots/blanks depict low/non-conserved amino acids.

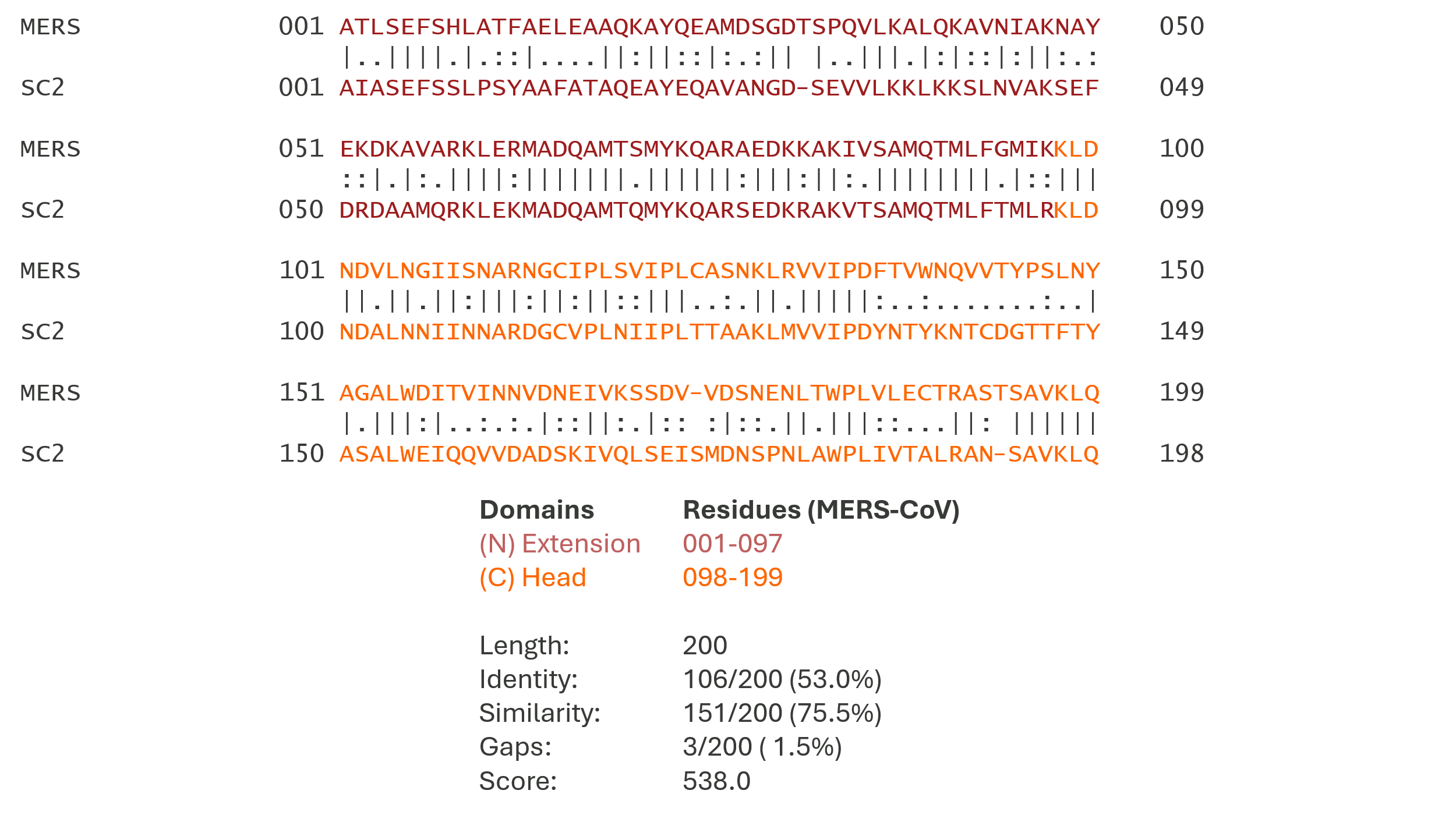

**Supplementary Figure 4. Sequence alignment of MERS-CoV and SARS-CoV-2 NSP8 using EMBOSS needle.** Coloured letters represent the domains; the N-terminal (Extension) domain in dark red and the C-terminal (Head) domain in orange.

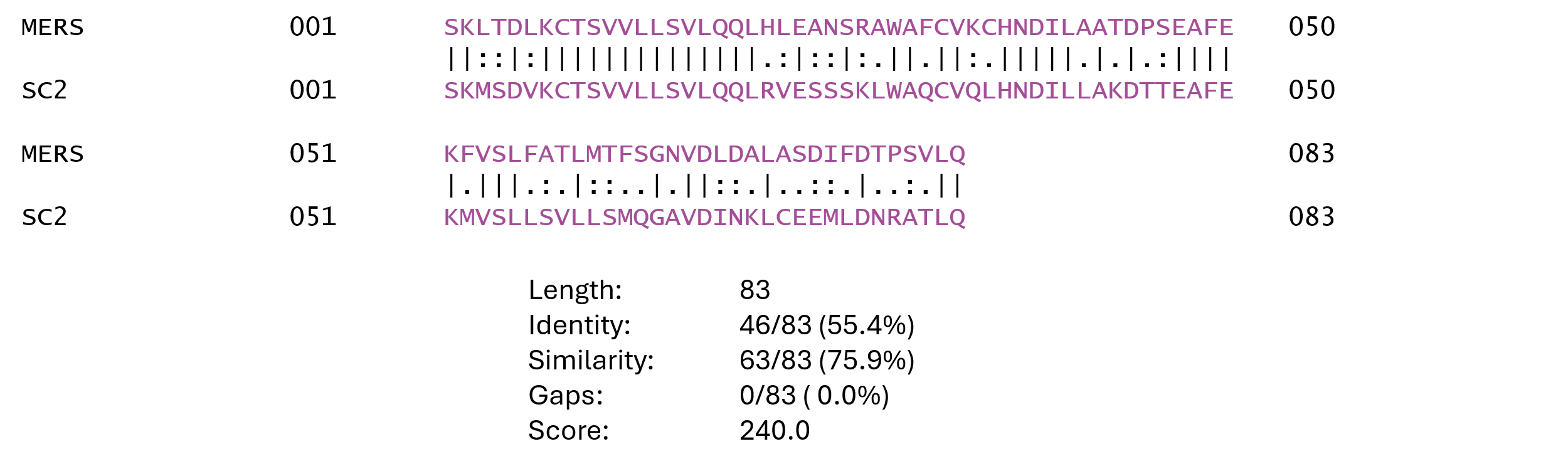

**Supplementary Figure 5. Sequence alignment of MERS and SARS-CoV-2 NSP7 using EMBOSS needle.** Vertical lines depict identical residues, colons depict semi-conserved residues and dot/blank depict low/non-conserved amino acids.

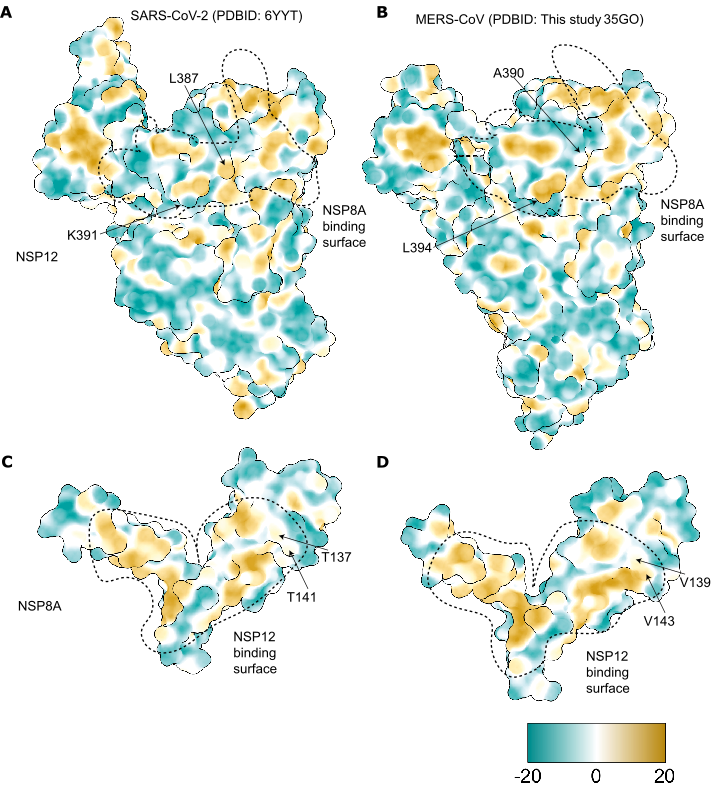

**Supplementary Figure 6. Comparison of hydrophobic interactions at the NSP12–NSP8a interface in SARS-CoV-2 and MERS-CoV RTCs.** Surface representations of NSP12 and NSP8A from the SARS-CoV-2 (PDB: 6YYT) and MERS-CoV (PDB: 35GO) RTC structures, coloured according to surface hydrophobicity (cyan, hydrophilic; yellow, hydrophobic). **A–B)** Comparison of the NSP8A-binding surface of NSP12 in SARS-CoV-2 (A) and MERS-CoV (B), highlighting differences in the distribution of hydrophobic regions surrounding the interaction interface. Residues K391 and L387 in SARS-CoV-2 NSP12 and their corresponding residues L394 and A390 in MERS-CoV NSP12 are indicated. **C–D)** Comparison of the NSP12-binding surface of NSP8A in SARS-CoV-2 (C) and MERS-CoV (D), highlighting differences in surface hydrophobicity and the positions of residues T137/T141 and V139/V143, respectively. Dashed outlines indicate the approximate interaction surfaces between NSP12 and NSP8A. Differences in the hydrophobic character and positioning of residues at the periphery of these interfaces may contribute to the asymmetric compatibility observed in cross-species NSP8 substitution experiments, potentially affecting the stability or geometry of NSP12–NSP8A interactions.

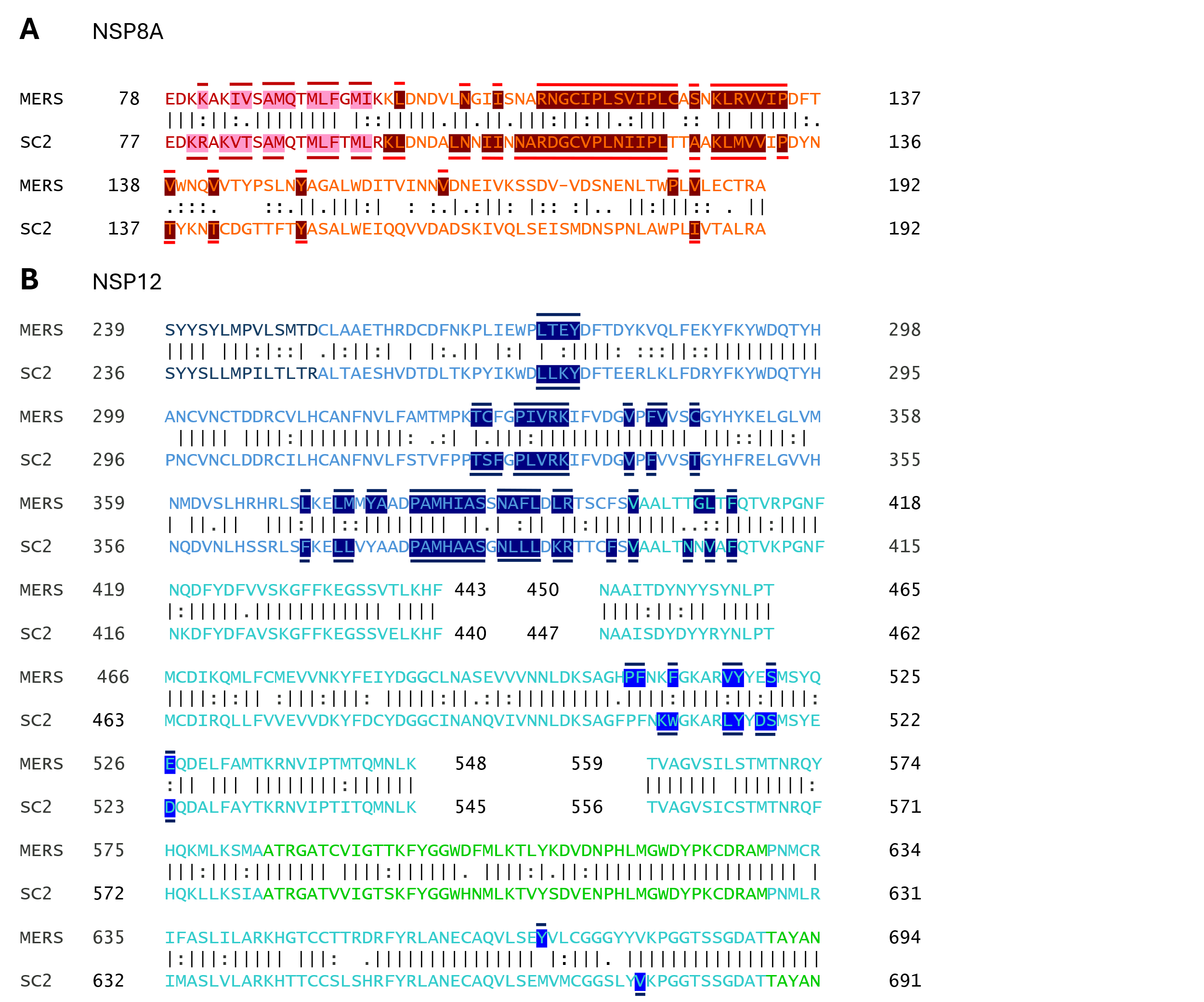

**Supplementary Figure 7.** **Sequence alignment of MERS-CoV and SARS-CoV-2 NSP8 and NSP12 interaction interfaces. A)** In the NSP 8 alignment, coloured letters represent the N-terminal extension domain (red) and C-terminal head domain (orange). Pink/dark red bars and highlights represent constituent components of the NSP8A-NSP12 interface. **B)** In the NSP12 alignment, dark blue, blue, teal and green colours represent the NiRAN, Interface, Fingers, and Palm domains, respectively. Dark blue bars and highlights represent components of the NSP12-8A interface. Interactions for MERS-CoV RdRp were derived from the generated structure featuring one copy of NSP8a and one of 12, whilst the SARS-CoV-2 interactions were derived from the PDB 6YYT structure (Hillen et al. 2020).

| RNA | Sequence 5' - 3' |
| --- | --- |
| Short primer | GCUAUGUGAGAU/Cy5/UAAGAAUU |
| Extended primer | GCUAUGUGAGAU/Cy5/UAAGAAUUAAAAAAAAAAAAAAAUUUUUUUU |
| Template | /Cy3/AAAAAAAAUUUUUUUUUUUUUUUAAUUCUUAAUCUCACAUAGC |
| -28 Template | /Cy3/AAAAAAAAUUUUUUUUUUUUUUUAAUUCUUAAUCUCACAUAGCGUAGCAUG |
| -28 primer | CAUGCUACGCUAUGUGAGAU/Cy5/UAAGAAUU |

**Supplementary Table 1.** Names and sequences of fluorescent RNAs used in the study.

|  | | **MERS_NSP12/8_RNA** | **MERS_NSP12/8x2/7_RNA** |
| --- | --- | --- | --- |
|  |  | EMD-100513 | EMD-100533 |
|  |  | PDB- 35GO/pdb_000035go | 35HQ/pdb_000035hq |
| **Data collection** | |  |  |
| Microscope | | Titan Krios(OPIC) | |
| Voltage (kV) | | 300 | |
| Detector | | Falcon 4i - SelectrisX | |
| Recording mode | | tiff | |
| Magnification | | 130,000 | |
| Movie/micrograph pixel size (Å) | | 0.932 | |
| Dose rate (e-/px/sec) | | 9.27 | |
| Number of frames per movie | | 40 | |
| Movie exposure time (s) | | 4.9 | |
| Total dose (e-/Å^2^) | | 50 | |
| Defocus range (um) | | 1.4 to 2.6 | |
| **EM data processing** | |  |  |
| Number of movies/micrographs | | 10, 942 | |
| Box size (px) | | 256 | |
| Particle number (After initial 3D Classes) | | 383, 291 | |
| Particle number (used in final map) | | 67,931 | 45,577 |
| Symmetry | | C1 | |
| Map resolution (Å, FSC 0.143) | | 3.27 | 3.4 |
| Local resolution range (Å, FSC 0.5) | | 2.9-30 | 3.0-30 |
| Map sharpening B-factor (Å^2^) | | -141 | -129 |
| **Model Building and Validation** | |  |  |
| Initial model used | | AlphaFold3 prediction | AlphaFold3 prediction |
| Model composition | |  |  |
| Non-hydrogen protein atoms | | 8,413 | 10,020 |
| Protein residues | | 994 | 1,202 |
| Nucleotides | | 20 | 20 |
| Ligand (Zn) | | 2 | 2 |
| B factors (Å^2^) - min/max/mean | |  | |
| Protein | | 16.8/177.4/71.4 | 11.9/227.8/88.3 |
| Nucleotide | | 44.9/152.2/87.1 | 41.1/153.4/91.5 |
| Ligand (Zn) | | 103.9/116.2/110.1 | 101.3/113.2/107.3 |
| RMSD from ideal | |  | |
| Bond length (Å) | | 0.004 | 0.004 |
| Bond angles (°) | | 0.575 | 0.588 |
| Validation |  |  | |
| Molprobity score | | 1.85 | 1.88 |
| Clashscore | | 6.34 | 8.41 |
| Rotamers outliers (%) | | 0 | 0 |
| FSC (0.5) model-vs-map | | 3.4 | 3.7 |
| CC model-vs-map (masked) | | 0.83 | 0.78 |
| Ramachandran plot | |  | |
| Favored (%) | | 91.5 | 93.4 |
| Allowed (%) | | 8.2 | 6.3 |
| Outliers (%) | | 0.3 | 0.3 |

**Supplementary Table 2.** Map and model statistics for cryo-EM.
